# Interplay of ERα binding and DNA methylation in the intron-2 determines the expression and estrogen regulation of Cystatin A in breast cancer cells

**DOI:** 10.1101/679043

**Authors:** Dixcy Jaba Sheeba John Mary, Girija Sikarwar, Ajay Kumar, Anil Mukund Limaye

## Abstract

Despite advances in early detection and treatment, invasion and metastasis of breast tumors remains a major hurdle. Cystatin A (CSTA, also called stefin A), an estrogen-regulated gene in breast cancer cells, is an inhibitor of cysteine cathepsins, and a purported tumor suppressor. Loss of CSTA expression in breast tumors evidently shifts the balance in favor of cysteine cathepsins, thereby promoting extracellular matrix remodeling, tumor invasion and metastasis. However, the underlying mechanism behind the loss of CSTA expression in breast tumors is not known. Here, we have analyzed CSTA expression, and methylation of upstream and intron-2 CpG sites within the CSTA locus in human breast cancer cell lines and breast tumors of the TCGA cohort. Results showed an inverse relationship between expression and methylation. Sequence analysis revealed a potential estrogen response element (ERE) in the intron-2. Analysis of ChIP-seq data (ERP000380) and our own ChIP experiments showed that 17β-estradiol (E2) enhanced ERα binding to this ERE in MCF-7 cells. This ERE was located amidst the differentially methylated intron-2 CpG sites, which provoked us to examine the possible conflict between estrogen-regulation of CSTA and DNA methylation in the intron-2. We analyzed the expression of CSTA and its regulation by estrogen in MDA-MB-231 and T47D cells subjected to global demethylation by 5-azacytidine (5-aza). 5-aza, not only enhanced CSTA expression in these cell lines but also restored estrogen-regulation of CSTA in these cells. Taken together, our results indicate that DNA methylation-dependent silencing could play a significant role in the loss of CSTA expression in breast tumors. The potential of DNA methylation as an indicator of CSTA expression or as a marker of tumor progression can be explored in future investigations. Furthermore, our results indicate the convergence of ERα-mediated estrogen regulation and DNA methylation in the intron-2, thereby offering a novel context to understand the role of estrogen-ERα signaling axis in breast tumor invasion and metastasis.

## 1. Introduction

Cysteine cathepsins are a major group of 11 lysosomal acid hydrolases in humans. The presence of cysteine in their active site distinguishes them from other lysosomal serine-or aspartic cathepsins (Khaket et al., 2019; Turk et al., 2012). Collectively, they hydrolyze proteins within and outside the cell (Turk et al., 2000). They are involved in various cellular processes, such as protein turnover, antigen presentation, apoptosis, and matrix remodeling (Fonović and Turk, 2014; Turk et al., 2000, 2012; Vizovišek et al., 2019). Overt or mislocalized expression of cysteine cathepsins is associated with diverse pathological conditions, including tumors and metastasis (Palermo and Joyce, 2008; Vidak et al., 2019). Cysteine cathepsins are regulated by compartmentalization, proteolytic cleavage (zymogen activation), pH and endogenous inhibitory proteins called cystatins (Turk et al., 2000, 2001; Verma et al., 2016). Cathepsins and cystatins are aberrantly expressed in a wide variety of tumors, including those of the breast (Pogorzelska et al., 2018). Altered expression of cysteine cathepsins or cystatins can tilt the homeostatic balance to favor extracellular matrix (ECM) remodeling, thereby promoting disease progression, tumor invasion and metastasis (Kolwijck et al., 2010; Paraoan et al., 2009; Yano et al., 2001).

Cystatin A (CSTA, also known as Stefin A), is a member of the Stefin group within the Cystatin superfamily (Keppler, 2006). It is an 11 kDa protein. It inhibits the activity of cathepsins -B, -H and -L (Rivenbark and Coleman, 2009), which are implicated in tumor development, progression and survival (Pogorzelska et al., 2018). It is abundantly expressed in the myoepithelial cells and plays a critical role in their ability to resist the malignant transformation of the luminal epithelium (Duivenvoorden et al., 2017; Jones et al., 2004). Estrogen reduces CSTA expression in MCF-7 breast cancer cells, via a mechanism that depends on the estrogen receptor α (ERα). Tamoxifen (a selective estrogen receptor modulator), fulvestrant (a selective estrogen receptor degrader), and ERα knockdown increase the expression of CSTA in MCF-7 breast cancer cells (John Mary et al., 2017). However, the precise role of ERα in the expression and estrogen-mediated regulation of CSTA in breast cancer cells is not understood.

The relationship between CSTA expression, and breast cancer progression and prognosis is confusing. In an immunohistochemical study on infiltrative breast carcinomas, Kuopio and co-workers found CSTA positivity in large tumors with higher mitotic activity. They also showed that CSTA positivity was associated with an increased risk of death (Kuopio et al., 1998). Lah and co-workers found lower CSTA mRNA and protein levels in breast carcinoma compared to their matched control in the majority of the 50 matched pairs under their study (Lah et al., 1992). Levicar and co-workers reported a 1.9 fold higher levels of CSTA in cytosols of primary invasive breast tumors compared to normal breast parenchyma. However, they did not find a significant association between CSTA and breast cancer prognostic factors (Levicar et al., 2002). Parker and co-workers analyzed CSTA expression in 142 primary breast tumors and found that CSTA expression not only correlated with disease-free survival but also decreased the risk of distant metastasis (Parker et al., 2008). Recently we analyzed the expression of CSTA in breast tumors from the TCGA cohort and reported significantly higher expression of CSTA mRNA in normal breast tissues compared with the primary breast tumors. High CSTA expression was associated with reduced overall (OS), relapse-free (RFS), and distant metastasis-free survival (DMFS). However, survival analysis of each subtype of breast tumors yielded interesting results. CSTA expression did not have any effect on survival in HER2+ and basal tumors. CSTA expression correlated with reduced OS and RFS in luminal A, whereas it correlated with prolonged OS and DMFS in luminal B (John Mary et al., 2017). Notwithstanding the conflicting results, these studies suggest a role for CSTA in breast cancer progression.

More recent studies on *in vivo* breast tumor metastasis models have provided valuable insights into the mechanistic role of CSTA in breast tumor progression. Parker and co-workers showed that CSTA expression and propensity to metastasize were inversely related in 4T1-derived cell lines, which exhibit varying degrees of metastasis in the syngeneic murine model. Furthermore, forced expression of CSTA in 4T1.2, a highly metastatic 4T1 line, reduced spontaneous metastasis to the bone (Parker et al., 2008). Using the same model, Withana et al showed that knockdown or selective inhibition of Cathepsin B can also reduce bone and lung metastasis (Withana et al., 2012). Taken together, these studies suggest that CSTA inhibits tumor invasion and metastasis by inhibiting Cathepsin B. Moreover, its tumor suppressor role is manifested in the tumor microenvironment. Loss of CSTA expression in primary tumors appears to be an important driving force towards metastatic progression. However, the mechanisms that lead to the loss of CSTA expression in breast tumors are not known.

Here, were have examined the relationship between CSTA expression and DNA methylation in specific regions within the CSTA locus in breast cancer cell lines, and breast tumors of the TCGA cohort. We show that CSTA expression has an inverse relationship with DNA methylation, suggesting that DNA methylation-dependent silencing could, at least in part, be responsible for the loss of CSTA expression in breast tumors. Furthermore, we found that estrogen regulation via ERα, and DNA methylation-dependent silencing converge in the intron-2 of CSTA.

## 2. Materials and methods

### 2.1. Plastic ware, chemicals and reagents

Cell culture plastic wares were purchased from Eppendorf (Hamburg, Germany). Cell culture media, fetal bovine serum (FBS), PowerUp SYBR Green PCR Master Mix, and High-Capacity cDNA Reverse Transcription Kit were from Invitrogen (CA, USA). Trypsin, penicillin, streptomycin, and charcoal-stripped FBS (csFBS) were from HiMedia (Mumbai, India). pMD20 vector was purchased from Clontech (CA, USA). Monoclonal β-Actin (AM4302) antibody and EpiJET Bisulfite Conversion Kit were purchased from Thermo Scientific (PA, USA). Monoclonal CSTA antibody (ab10442) was from Abcam (Cambridge, UK) and polyclonal ERα antibody (sc-543) was purchased from Santa Cruz Biotechnology (CA, USA). Progesterone Receptor (PR) antibody (8757S) was from Cell Signaling Technology (Massachusetts, USA). 5-azacytidine (5-aza) was procured from MP Biomedicals (Solon, USA). 17β-estradiol (E2) were purchased from Sigma Aldrich (MO, USA). Routine laboratory buffers, solvents and salts were from Merck (Mumbai, India) or SRL (Mumbai, India).

### 2.2. Cell lines and cell culture

MCF-7 cells were grown in Dulbecco’s Modified Eagle’s Medium (DMEM) with phenol red. T47D, ZR-75-1, MDA-MB-231, and MDA-MB-453 were grown in Rosewell Park Memorial Institute-1640 medium (RPMI-1640) with phenol red. These media for routine culture were supplemented with 10% (v/v) heat-inactivated FBS, 100 units/ml penicillin and 100 μg/ml streptomycin (M1 medium) in a humidified 37 °C incubator with 5% CO_2_. For experiments involving estrogen treatment, phenol red-free DMEM/F12 or RPMI-1640 media supplemented with heat-inactivated csFBS, 100 units/ml penicillin and 100 μg/ml streptomycin (M2 medium) were used.

### 2.3. Treatment protocols

#### 2.3.1. Global demethylation using 5-aza

Cells were seeded in M1 medium. After 24 h, the cells were treated with fresh M1 medium containing 10 µM 5-aza for 5 days. Fresh M1 medium with 5-aza was replenished every 48 h. The cells treated with DMSO (vehicle) served as control.

#### 2.3.2. E2 treatment on cells subjected to global demethylation

MDA-MB-231 or T47D cells were treated with 5-aza as described above, followed by incubation in M2 medium for 4 h. Thereafter, the cells were treated with 10 nM E2 or ethanol (vehicle) in M2 medium. After 24 h the cells were harvested for isolation of total RNA or protein.

#### 2.3.3. Treatment of MCF-7 cells with E2 for chromatin immunoprecipitation (ChIP) assay

MCF-7 cells were seeded in 100 mm dishes in M1 medium. When the cells were 70 % confluent, the spent medium was replaced with M2 medium and incubated for 24 h. Thereafter, the cells were treated with fresh M2 medium containing 10 nM E2 or ethanol (vehicle). After 24 h, the cells were harvested for ChIP assay.

### 2.4. Routine reverse transcription-polymerase chain reaction (RT-PCR) and quantitative RT-PCR (qRT-PCR)

Total RNA was isolated, reverse transcribed and the resultant cDNA was subjected to routine RT-PCR and qRT-PCR analyses with gene-specific primers (Supplementary table 1) for CSTA, ERα and cyclophilin A (CycA) as previously described (John Mary et al., 2017). CycA served as an internal control

**Table 1:** Correlation between CSTA expression and methylation in CSTA locus in breast tumors of the TCGA cohort

| Probe ID | Spearman's $\rho$ | $p$-value |
| --- | --- | --- |
| cg14664412 | -0.5482637 | < 0.0001 |
| cg18618429 | -0.5683092 | < 0.0001 |
| cg21932814 | -0.4717445 | < 0.0001 |
| cg26928972 | -0.5154166 | < 0.0001 |

### 2.5. Western blotting

Total protein was isolated from breast cancer cells with Laemmli sample buffer (Laemmli, 1970) or from organic phase of TRIzol lysates and then quantified by TCA method (Karlsson et al., 1994) or Lowry’s method (Lowry et al., 1951) respectively. Protein samples (30 µg) were resolved by 12% PAGE, transferred to 0.22 µ nitrocellulose membrane, and blocked with 1% gelatin in TBST for 2 h. Blots were probed with CSTA, ERα, PR, β-actin, or histone H3 antibodies, overnight, in TBST containing 0.1% gelatin. The blots were washed with TBST (6 × 5 min). Blots were then probed with anti-rabbit HRP-conjugated secondary antibody for 1 h, washed with TBST (6 × 5 min) and developed using Clarity Western ECL Substrate (Bio-Rad, California, US). Images were captured with ChemiDoc XRS+ system (Bio-Rad, California, US).

### 2.6. TCGA-BRCA data analysis

Methylation (generated with Illumina Infinium® Human Methylation 450K BeadChip array) and CSTA expression (RNAseq) data were retrieved from TCGA database (Koboldt et al., 2012) using the UCSC Xena Browser (Goldman et al., 2019). CSTA expression data were available for 1218 samples. Out of these, 873 samples (85 normal and 788 tumors) had both expression and methylation data. Only the tumor samples were used for analysis. Data were processed in MS Excel. Correlation between methylation (beta values) and CSTA expression (log_2_RPKM+1) was assessed by Spearman’s rank correlation test. Further, tumors were classified into hyper-methylated and hypo-methylated based on a threshold beta-value of 0.3. CSTA expression in hyper-methylated and hypo-methylated tumors was represented as box plots. The difference in CSTA expression in these two groups was analysed by Welch two-sample *t*-test.

### 2.7. Bisulfite sequencing

Genomic DNA (gDNA) was isolated from breast cancer cell lines. Two µg of gDNA was bisulfite converted and purified with EpiJET Bisulfite Conversion Kit. 50 ng of converted gDNA was used for PCR with specific primers (Supplementary table 1) that amplified Region 1 and Region 2, which encompassed few CpG dinucleotides of the upstream region and intron-2, respectively. The primers were designed to amplify only the bisulfite converted DNA. Importantly, we ensured that the priming region did not contain any CpG dinucleotides. The PCR products were gel purified and cloned in pMD20 vector and sequenced. The inserts of 12 −13 independent clones per cell line were sequenced. The sequencing results were analyzed to determine methylated and unmethylated CpG sites and represented as lollipop plots. The proportion of CpGs methylated in Region 1 or Region 2 for each cell line was determined.

### 2.8. ChIP

Control and E2-treated MCF-7 cells were fixed with formaldehyde (0.75%) for 10 minutes. The reaction was stopped by 125 mM glycine for 10 minutes. Cells were washed and scraped with ice-cold DPBS, pelleted, lysed with lysis buffer (50 mM HEPES pH 7.5, 150 mM KCl, 1 mM EDTA, 10% glycerol, 0.1% NP-40), and sonicated. Lysates were clarified by centrifugation, and supernatants containing chromatin were collected. Chromatin samples were pre-cleared with Protein G plus-Agarose beads precoated with BSA and herring sperm DNA for 2 h. 5% of the pre-cleared chromatin samples were separated as input. The remaining portions were incubated with ERα antibody or rabbit IgG antibody for 2 h. Samples were immuno-precipitated by incubating with 20 μl of coated beads for 2 h, followed by centrifugation. Immunoprecipitates were washed with a series of wash buffers (Shivaswamy and Iyer, 2007) and eluted in 300 µl elution buffer containing proteinase-K for 2 h at 55°C, and overnight incubation at 65°C. Chromatin was column purified and ERα occupancy was assessed by PCR with primers specific to pS2 (positive control) or Region 2 of CSTA locus (Supplementary table 1).

### 2.9. ChIP-Seq analysis

Raw data of Chip-Seq experiments were retrieved from Sequence Read Archival (SRA) and analyzed using Galaxy, web-based platform (Afgan et al., 2016). Chip-Seq data (SRA accession ID: ERP000380) of chromatin samples from MCF-7 cells treated with E2 (ID:ERR022026), tamoxifen (ID:ERR022027) or vehicle (ID:ERR022025) and immuno-precipitated with ERα was chosen for this study. IgG control ChIP-seq data was used as the negative control. Using FASTQC tool (Andrews, 2010), the quality of input reads were assessed. After converting the quality score to sanger quality type by FASTQ Groomer (Blankenberg et al., 2010), reads were mapped to reference human genome (hg19) using “Map with Bowtie for Illumina” tool (Langmead et al., 2009). Unmapped reads were discarded by “Filter SAM or BAM, output SAM or BAM” tool (Li et al., 2009). Genomic regions with enriched sequencing reads were identified by MACS (Model-based analysis of ChIP-Seq) tool (Zhang et al., 2008). Resultant Wig files were converted to bigWig files using “Wig/BedGraph-to-bigWig” tool and the peaks representing ERα occupancy were visualized using UCSC genome browser (Kent et al., 2002).

### 2.10. Statistical analysis

Bisulfite sequencing data were analyzed for statistical difference in proportions of methylated CpGs for every pair of cell lines. The total number of CpGs sampled for each cell lines was greater than 30. We assumed that methylation at each CpG site was independent of the adjacent CpGs. Under null hypothesis, the mean of the sampling distribution of the difference in proportions follows a standard normal (z) distribution. The test statistic *d* was calculated as

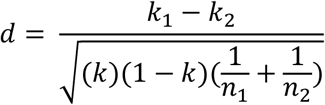

where, *k*_*1*_*-k*_*2*_ is the observed difference in proportion in a pair of cell lines, *k* is the proportion for the pooled data of the pair of cell lines, and *n*_*1*_and *n*_*2*_are the number of CpGs sampled. The probability of *d*was obtained from z distribution. The p-value obtained for each pair of cell lines was subjected to Bonferroni correction.

Correlation between methylation score and CSTA expression was analyzed as described in section 2.6. All other data were analyzed by ANOVA followed by Tukey’s HSD, or Welch two-sample *t*-test using R-statistical package.

## 3. Results

### 3.1. Differential expression of CSTA in breast cancer cell lines

We analyzed CSTA mRNA and protein expression in a panel of breast cancer cell lines using RT-PCR and western blotting techniques, respectively. CSTA mRNA expression was highest in ZR-75-1 cells, followed by MCF-7. T47D cells expressed low but detectable levels of CSTA mRNA. CSTA mRNA expression in MDA-MB-231 and MDA-MB-453 was undetectable (Fig. 1A). CSTA protein expression matched the mRNA expression in these cell lines (Fig. 1B). T47D cells expressed the highest levels of ERα, followed by MCF7. ZR-75-1 cells expressed low but detectable levels of ERα. ERα expression was undetectable in the remaining cell lines (Fig. 1A, B).

**Fig 1.**
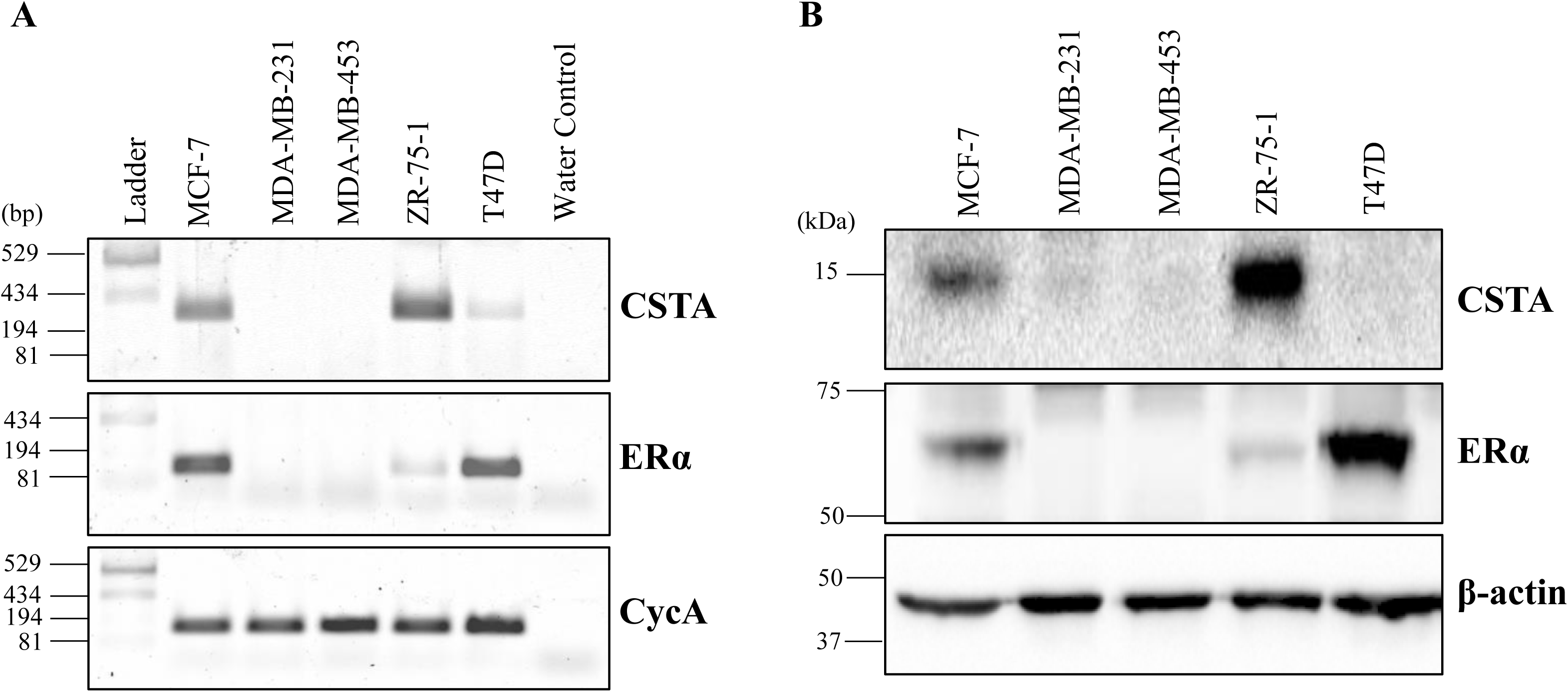
Expression of CSTA in breast cancer cell lines. A. Total RNA was isolated from the indicated breast cancer cell lines and subjected to RT-PCR analysis using primers specific for CSTA, ERα, and CycA. CycA served as an internal control. B. Total protein was isolated from the indicated breast cancer cell lines, and subjected to western blot analysis using primary antibodies specific to CSTA, ERα, and β-actin. β-actin served as an internal control.

### 3.2. 5-aza induces CSTA expression in MDA-MB-231 cells

Treatment with 5-aza, a DNA methyltransferase 1 (DNMT1) inhibitor, causes genome-wide demethylation. We tested the effect of 5-aza on CSTA expression in MDA-MB-231 cells. As shown in Fig. 2, 5-aza treatment induced CSTA mRNA expression. This suggested that low or absence of CSTA expression in breast cancer cells could be a result of DNA methylation in the CSTA locus.

**Fig. 2.**
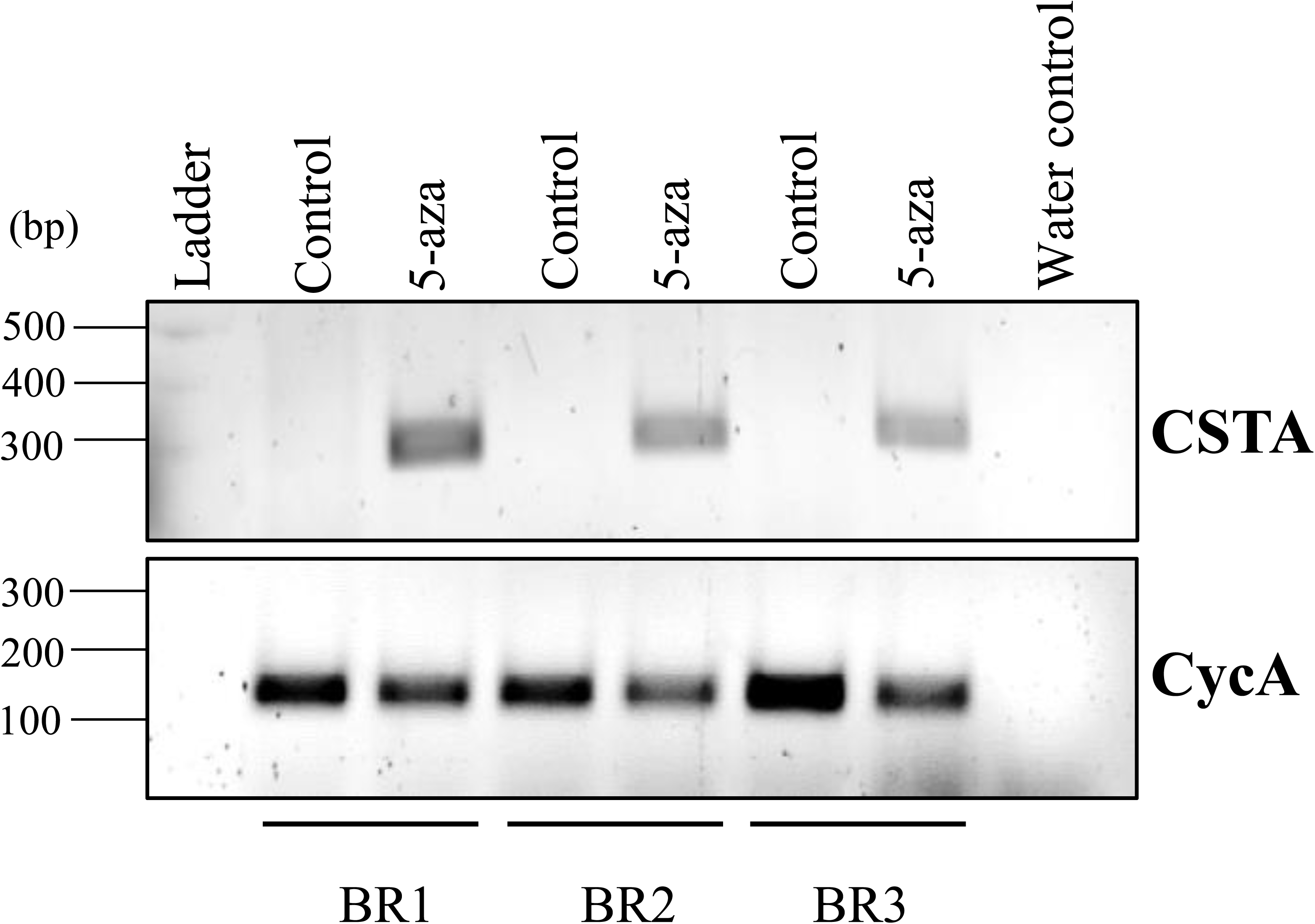
5-aza induces the expression of CSTA mRNA in MDA-MB-231 cells. MDA-MB-231 cells were treated with DMSO (vehicle control) or 5-aza for 5 days in M1 medium. Total RNA was isolated and reverse transcribed. The expression of CSTA was analysed by routine RT-PCR. BR1, BR2, and BR3 are three biological replicates. Each biological replicate comprised of one dish each for DMSO and 5-aza treated cells.

### 3.3. *In silico* analysis of DNA methylation in the CSTA locus

CSTA does not have CpG islands (Rivenbark and Coleman, 2009). Methylation data obtained from 450K bead chip arrays (Encode project) was analyzed to ascertain the status of methylation in five CpGs in the CSTA locus. The locations of these CpGs is indicated by the colored vertical lines in Fig. 3A. One of these mapped to the intron-2. Among the remaining (referred to as the upstream CpGs), two mapped to the exon-1, and two were present in the upstream region close to the transcription start site. In MCF-7 cells, the four upstream CpGs appeared methylated. In T47D cells, out of the four upstream CpGs, one was methylated, one was unmethylated and the remaining two were partially methylated. The intron-2 CpG appeared unmethylated in MCF-7, but partially methylated in T47D cells (Fig. 3A). The *in silico* analysis indicated that methylation in the intron-2 CpG had an inverse relationship with CSTA expression in the two cell lines.

**Fig. 3.**
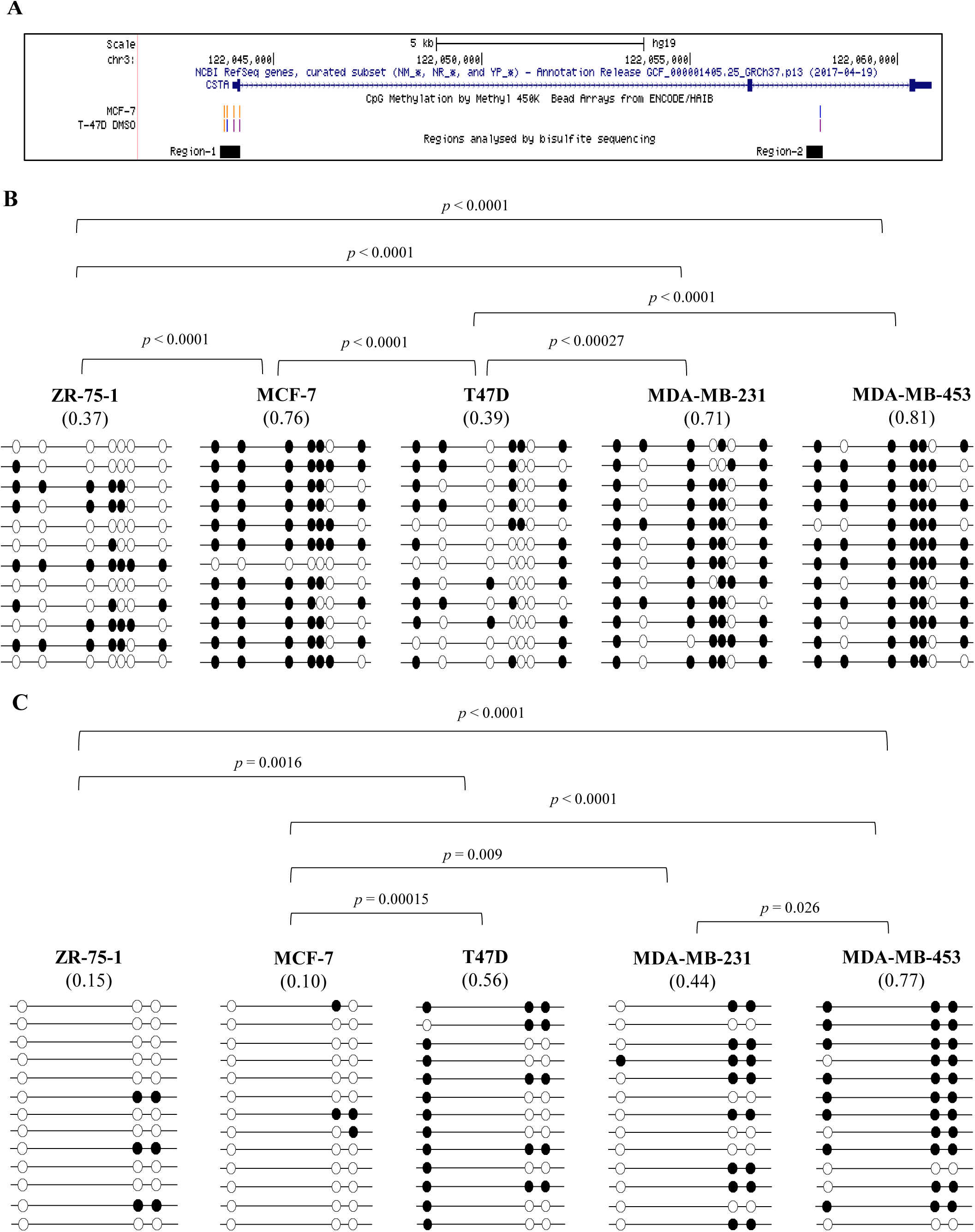
Differential methylation of upstream and intron-2 CpG sites of the CSTA locus in breast cancer cell lines. A. Snapshot from UCSC genome browser displaying the location of Methylation 450K BeadChip array probes with respect to the CSTA locus. The colored vertical lines along the MCF-7 and T47D tracks indicate the extent of methylation of the CpG sites; orange = methylated (beta value >= 0.6), purple = partially methylated (0.2 < beta value < 0.6), bright blue = unmethylated (0 < beta value <= 0.2). Black solid rectangles labeled as Region 1 and Region 2 indicate the regions analyzed by bisulfite sequencing. B, C. Bisulfite sequencing of Region 1 and Region 2, respectively. gDNA samples isolated from the indicated breast cancer cell lines were bisulfite converted and used for PCR reactions with primers specific to Region 1 and Region 2. The PCR amplified products were cloned in TA vector and sequenced. The inserts from 12 or 13 independent TA clones per cell line were analyzed for methylated and unmethylated CpG sites in Region 1 and Region 2, respectively. The methylation pattern is represented by lollipop plots. Filled circles represent methylated CpGs, and open circles represent unmethylated CpGs. The numbers below each cell line indicate the proportion of methylated CpGs. Two proportions from each pair of cell lines were tested for significant difference as described in materials and methods. The indicated *p*-values are adjusted *p*-values obtained following Bonferroni correction. which are indicated only for the pair of cell lines with significant difference in the proportion of methylated CpGs.

### 3.4. Bisulfite sequencing of upstream and intron-2 regions in the CSTA locus

We isolated gDNA from the panel of breast cancer cells indicated in Fig. 1. They were subjected to bisulfite sequencing analysis of Region 1 and Region 2 (Fig. 3A), which harbor the CpG dinucleotides interrogated with the Encode data. Region 1 and Region 2 have a total of 7 and 3 CpG dinucleotides respectively. 84 CpGs were interrogated across 12 independent TA clones in the Region 1. 39 CpGs were interrogated across 13 independent clones in Region 2. The lollipop models for methylated and unmethylated CpG sites in Regions −1 and −2 are shown in Fig. 3B and 3C, respectively. We determined the proportion of methylated CpGs in each region in the cell lines. In the Region 1, MDA-MB-453 cells showed the highest proportion of methylated CpGs, followed by MCF-7, MDA-MB-231, T47D and ZR-75-1, in the decreasing order. In the Region 2, MDA-MB-453 cell showed the highest proportion of methylated CpGs, followed by T47D, MDA-MB-231, ZR-75-1 and MCF-7 cells, in the decreasing order. There were significant differences in the proportions of methylated CpGs in both the regions in the indicated pair of cell lines (Fig. 3B and 3C). Generally, methylation in both regions appeared to be inversely related to CSTA expression in the breast cancer cell lines.

### 3.5. Methylation of the upstream CpGs correlates inversely with CSTA expression in the breast tumors

CSTA expression (RNASeq; log_2_(RPKM+1)) and methylation data (generated with Illumina Infinium® Human Methylation 450K BeadChip array) for the primary breast tumors in the TCGA breast cancer dataset were accessed using the UCSC Xena Browser (Goldman et al., 2019). Methylation data were available for 4 probes in the CSTA locus. These probes correspond to the four CpG sites in the Region 1 shown in Fig. 3A. We first performed a probe-wise analysis of the correlation between methylation (beta value) and CSTA expression in primary tumors. Methylation at each of the 4 CpG sites inversely correlated with CSTA expression (Table 1). The primary tumors were divided into two groups, namely hypo-and hyper-methylated, using a beta value of 0.3 as a cut-off. CSTA expression in hypo-methylated tumors was significantly higher than those in hyper-methylated tumors (Fig. 4A-D). We generated a composite methylation score for each sample by averaging the beta values for all the probes. The composite methylation score correlated inversely with CSTA expression (ρ = −0.582, *p* < 0.0001, Fig. 4E). Furthermore, when the tumors were divided into hypo-and hyper-methylated groups, based on a cut-off composite methylation score of 0.3, the hypo-methylated tumors showed significantly higher expression of CSTA compared to hyper-methylated tumors (Fig. 4F).

**Fig. 4.**
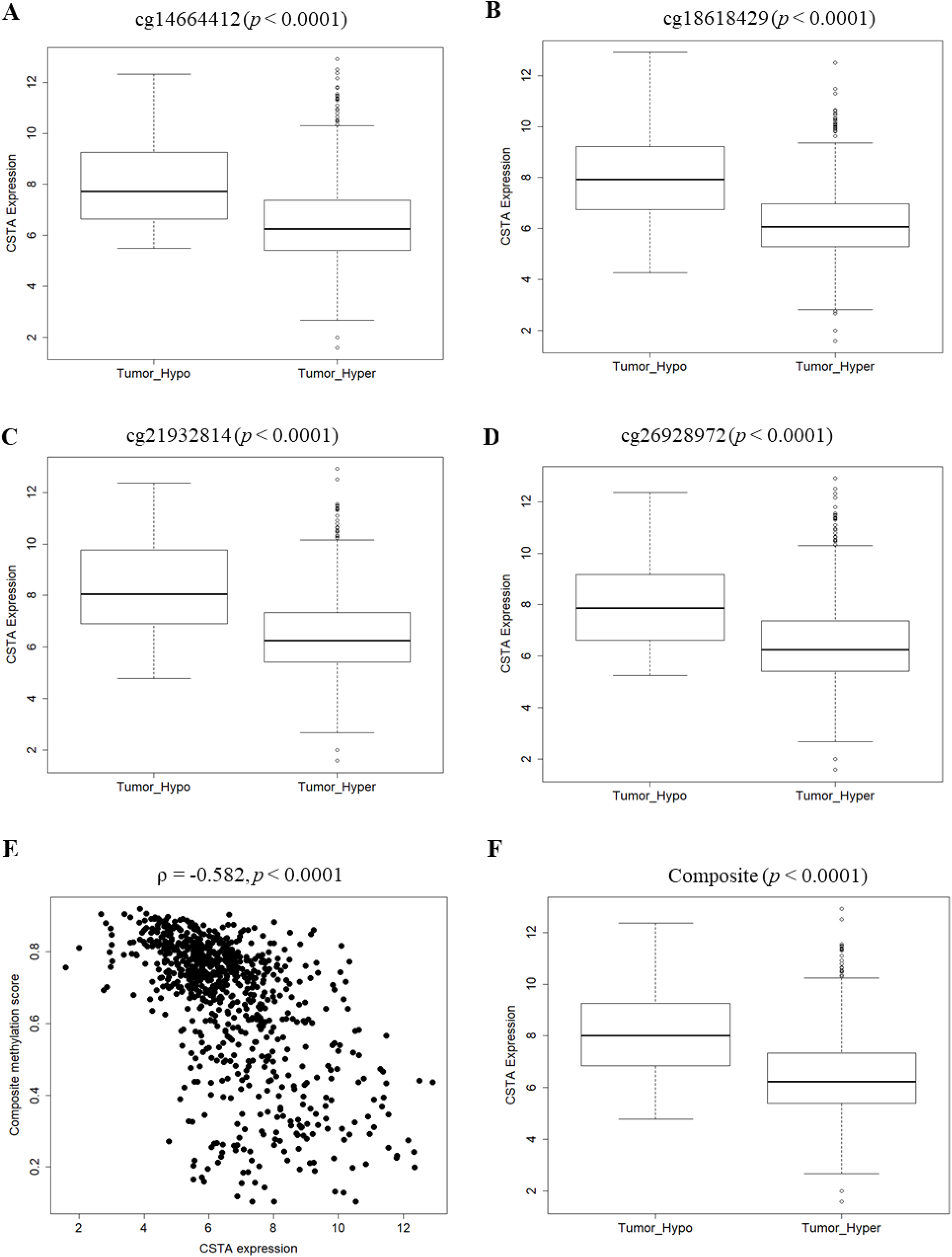
Inverse correlation between CpG methylation and CSTA expression in breast tumors of the TCGA cohort. A-D. Probe-wise analysis of the correlation between methylation and CSTA expression. The tumors were segregated into hypo-methylated or hyper-methylated groups based on the thrershold beta value of 0.3 for each probe. The distribution of CSTA expression in hypo-methylated and hyper-methylated tumors are represented as box plots. E-F. A composite methylation score, which is the average beta value of all the probes, was determined for each tumor sample. The scatter plot of the composite methylation score versus CSTA expression is shown in E. The tumors were segregated into hypo-methylated or hyper-methylated groups based on the thrershold composite score of 0.3. The distribution of CSTA expression in hypo-methylated and hyper-methylated tumors is represented as a box plot (F). The difference in mean CSTA expression in hypo-methylated and hyper-methylated tumors was analysed by Welch two-sample *t*-tests.

### 3.6. Estrogen enhances ERα occupancy in the intron-2 region of CSTA in MCF-7 cells

Analysis of the CSTA locus using JASPAR (Sandelin et al., 2004) revealed a potential estrogen response element (ERE) in the intron-2 region. This ERE was located amidst the CpG sites analyzed in Region 2 (Fig. 5A). Chip-Seq data of chromatin samples from vehicle-, E2-or tamoxifen-treated MCF-7 cells, which were immunoprecipitated with ERα-specific antibody, were analyzed using Galaxy. The results were viewed in the UCSC genome browser. A robust peak of ERα occupancy was observed in the intron-2 region of CSTA in E2-treated MCF-7 cells (indicated by the red rectangle, Fig. 5B). This peak was diminished or negligible in tamoxifen or vehicle treated MCF-7 cells. Notably, the peak of ERα binding overlapped with ERE predicfted by JASPAR. To validate these observations, we performed ChIP experiments on ethanol (vehicle) or E2-treated MCF-7 cells using an ERα specific antibody. Enrichment of the ERE containing sequence in the pS2 locus (positive control) following E2 treatment validated the ChIP protocol. As shown in Fig. 5C, the Region-2 was enriched in immunoprecipitated chromatin samples from E2-treated MCF-7 cells.

**Fig. 5.**
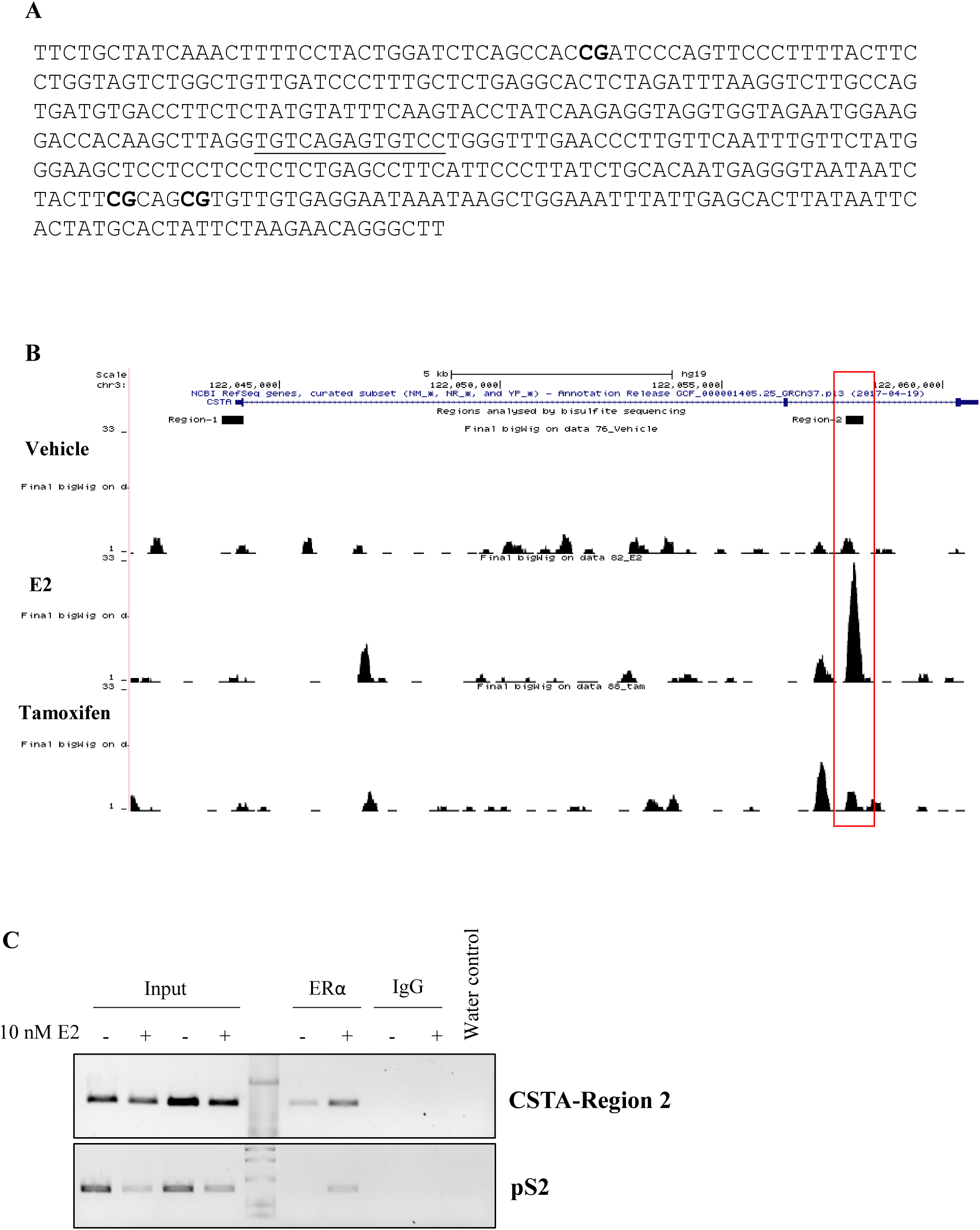
Estrogen enhances ERα occupancy in the intron-2 region of CSTA in MCF-7 cells. A. The location of ERE predicted by JASPAR in the Region 2. The underlined sequence is the ERE. The CpG sites analysed by bisulfite sequencing are indicated by bold letters. B. A snapshot of the UCSC genome browser displaying the results of ChIP-Seq analysis. Fastq files of ChIP-Seq data (Project ID: ERP000380) were retrieved from SRA repository and analysed in Galaxy. Note that E2 treatment increases the ERα occupancy in the intron-2 in a region that overlaps with Region 2 (red rectangle). C. The result of ChIP experiment. MCF-7 cells were treated with E2 for 24 h. Cross-linked chromatin samples from the treated and control cells were fragmented and immunoprecipitated with ERα or IgG specific antibodies. Immunoprecipitated DNA was reverse cross-linked, purified and subjected to PCR analysis using primers flanking the intron-2 ERE. pS2, a known E2 induced gene served as a positive control. Data shown are representative of three independent experiments.

### 3.7. Global demethylation restores estrogen regulation of CSTA in MDA-MB231 and T47D cells

In MDA-MB-231 cells, DNA methylation-dependent silencing is the underlying mechanism for the loss of ERα (Lapidus et al., 1998) and CSTA (this study) expression. We examined ERα and CSTA expression in 5-aza-untreated or -pretreated MDA-MB-231 cells, which were stimulated with vehicle or 10 nM E2. As expected, in 5-aza-untreated cells, ERα protein expression was not detectable after vehicle or E2 treatment (Fig. 6A, lanes 1 and 2). There was no significant difference in CSTA mRNA expression (Fig. 6C, bars 1 and 2, ANOVA followed by Tukey’s HSD). On the other hand, in 5-aza-pretreated cells, an immunoreactive protein was detected on western blots with ERα specific antibody (Fig. 6A, lanes 3 and 4). This immunoreactive protein had a higher molecular mass compared to the expected 66 kDa for ERα. Notwithstanding this discrepancy, induction of PR, and further enhancement of its expression with E2 confirmed the generation of a functional ERα in 5-aza pretreated cells (Fig. 6A, B, lanes 3 and 4). 5-aza significantly induced CSTA mRNA expression in MDA-MB-231 cells (Fig. 6C, bars 1 and 3; ANOVA followed by Tukey’s HSD). E2 suppressed the 5-aza induced levels of CSTA mRNA, although the difference was not statistically significant when analyzed by ANOVA. However, the levels of CSTA mRNA in 5-aza pre-treated cells with and without E2 treatment were significantly different when analyzed by the Welch two-sample *t*-testin (Fig. 6C, bars 3 and 4, *p* = 0.0098). Western blots failed to demonstrate CSTA protein in MDA-MB-231 cells.

**Fig. 6.**
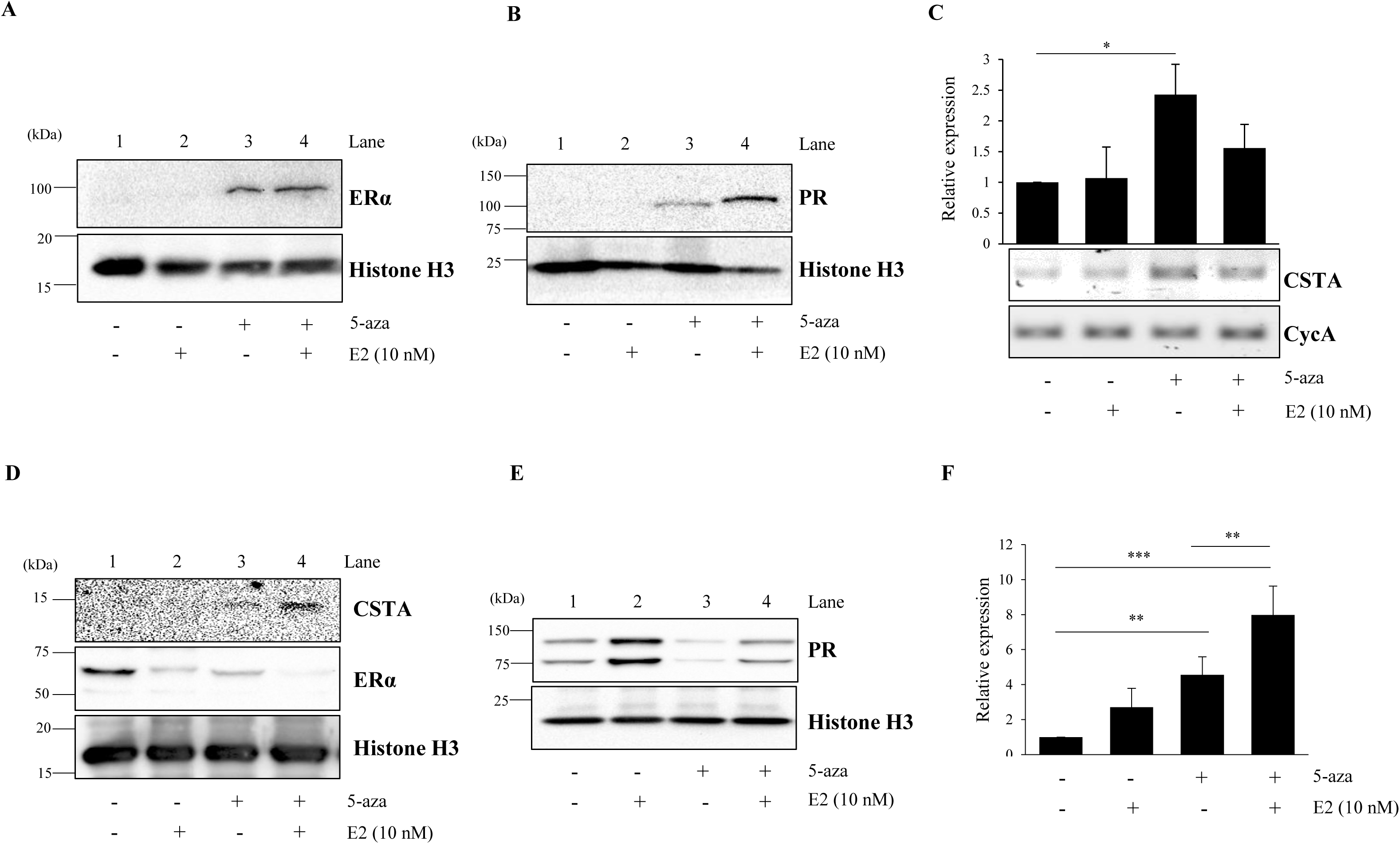
Global demethylation restores estrogen regulation of CSTA. MDA-MB-231 (A-C) or T47D (D-F) cells were subjected to global demethylation using 5-aza. The cells were then stimulated with 10 nM E2 or ethanol (vehicle) for 24 h. CSTA expression was analyzed by western blotting (A, B, D, E) and RT-PCR (C) or qRT-PCR (F). CycA was used as internal control in RT-PCR and qRT-PCR analyses. Histone H3 served as an internal control in western blots. Relative CSTA mRNA expression data are represented as bar graphs. The expression in the control samples (without 5-aza and E2 treatments) were set to 1 and the expression in the other treatment groups were expressed relative to the control. Bars represents mean relative expression (± S.D.) Data were analyzed by ANOVA followed by Tukey’s HSD. n = 3 biological replicates for MDA-MB-231 and n = 4 biological replicates for T47D. Each biological replicate comprises of one dish each for control and treated cells. \**p* < 0.05, ** *p* < 0.01, \*\*\**p* < 0.001.

We applied the same experimental design to ERα positive T47D cells, which express a very low or undetectable level of CSTA. Without 5-aza pretreatment, T47D cells treated with E2 showed decreased levels of ERα protein and increased levels of PR, as expected (Fig. 6D, E, lanes 1 and 2) (Al-Dhaheri et al., 2006; Wormke et al., 2000). There was no observable effect on CSTA protein (Fig. 6D, lanes 1 and 2). However, we observed an increase in CSTA mRNA (Fig. 6F, bars 1 and 2; ANOVA followed by Tukey’s HSD), although the increase was not statistically significant. 5-aza pre-treatment alone caused a decrease in ERα protein in T47D cells (Fig. 6D, lanes 1 and 3) in a manner similar to that reported in MCF-7 cells (Pryzbylkowski et al., 2008). This also led to increased CSTA (Fig. 6D, lanes 1 and 3) and decreased PR protein expression (Fig. 6E, lanes 1 and 3). E2 induction of PR in 5-aza pre-treated T47D cells showed that ERα was functional (Fig. 6E, lanes 3 and 4). E2 not only enhanced CSTA protein expression (Fig. 6D, lanes 3 and 4), but also significantly enhanced CSTA mRNA in 5-aza pre-treated cells (Fig. 6F, bars 1, 3 and 4; ANOVA followed by Tukey’s HSD).

## 4. Discussion

There were two motivating reasons behind our study of the relationship between CSTA gene expression and methylation in breast cancer. Firstly, CSTA is a proposed tumor suppressor (Calkins and Sloane, 1995; Duivenvoorden et al., 2017), and methylation-dependent silencing of tumor suppressors are well known (Kazanets et al., 2016). Recently, Ma and co-workers showed the association between the loss of CSTA expression in lung cancer cell lines and partial methylation of the CpG dinucleotides in the promoter and exon 1 regions (Ma et al., 2018). Secondly, Cystatin M, which is also a reversible inhibitor of Cathepsins -B and -L, is silenced due to methylation of the proximal promoter CpG island in breast cancer cell lines and primary invasive breast tumors (Ai et al., 2006; Rivenbark et al., 2006b, 2006a; Schagdarsurengin et al., 2007). Our analysis focused on CpG methylation on two regions of the CSTA locus. This is the first study to demonstrate the inverse relationship between CSTA expression and methylation in the context of breast cancer. The CSTA locus lacks CpG islands. There are examples of DNA methylation-dependent regulation of CpG island-less genes (Arapshian et al., 2000; Deng et al., 2004; Graff et al., 2000; Sathyanarayana et al., 2004, 2003a, 2003b, 2003c). Hence, our results are not surprising.

The conclusions of this study are drawn from results obtained through three different approaches; a) analysis of CSTA expression and methylation data from the TCGA breast cancer cohort, b) examination of DNA methylation data (ENCODE project) for the CSTA locus, and c) bisulfite sequencing of DNA isolated from breast cancer cell lines, which express differential levels of CSTA. TCGA methylation data was generated using the Infinium methylation 450K bead chip arrays, which do not have probes to interrogate all CpG sites in the CSTA locus. Due to this limitation, no conclusion could be drawn regarding the correlation between CSTA expression and methylation of the intron-2 CpGs. Nevertheless, an inverse correlation between CSTA expression and methylation in the upstream CpG sites was observed (Fig. 4E). The ENCODE project data corresponding to MCF-7 and T47D cells, and our bisulfite sequencing results clearly demonstrated that CSTA expression, and methylation in the intron-2 CpGs, are inversely correlated. Collectively, these data provide compelling evidence in favor of DNA methylation-dependent silencing of CSTA in breast cancer cells. Altered Cathepsin B: CSTA ratio in breast tumors is reported. It also correlates with disease prognosis (Lah et al., 1997, 1992; Levicar et al., 2002). Ablation or inhibition of Cathepsin B also inhibits spine and lung metastasis in the animal model (Withana et al., 2012). We therefore, propose that DNA methylation-mediated silencing of CSTA in primary breast tumors tips the CTSB/CSTA balance in favor of CTSB, which in turn facilitates tumor invasion and metastasis. A detailed study on the correlation of DNA methylation in the CSTA locus, and disease progression, treatment outcome and survival, may uncover its potential as a prognostic marker.

Our previous analysis of CSTA and ERα mRNA expression levels in breast tumors of the TCGA cohort had revealed an inverse correlation. This is consistent with the observed induction in CSTA mRNA in MCF-7 cells following ERα knockdown (John Mary et al., 2017). The inverse relationship between CSTA and ERα expression is reiterated in the ERα positive breast cancer cell lines used in this study. ZR-75-1 which has the highest expression of CSTA has the least ERα expression, whereas T47D, which has the least expression of CSTA has the highest expression of ERα. CSTA and ERα expression levels in MCF-7 cells are in between these two extremes. In the previous study, we also showed that estrogen suppressed CSTA expression in MCF-7 cells via ERα (John Mary et al., 2017). However, the mechanistic role of ERα was not known. The prediction of an ERE by JASPAR, the peak of ERα binding revealed by ChIPseq data, and our own validation of increased ERα occupancy in the intron-2 following estrogen treatment of MCF-7 cells suggest that estrogen suppresses CSTA expression at the level of transcription. The precise events post -ERα binding that leads to transcriptional shut-off are worth addressing in future investigations. However, the mechanism of estrogen-mediated regulation of CSTA is more complex. Estrogen does not produce similar effects on CSTA expression in ERα positive breast cancer cell lines. Estrogen suppresses CSTA expression in ZR-75-1 cells. However, the extent of suppression is much lower than that observed in MCF-7 cells. In T47D cells, estrogen does not modulate CSTA expression. The study offers no insight into the reasons behind differential responses of the breast cancer cell lines to estrogen stimulation. However, it highlights the role of intron-2 methylation in estrogen regulation of CSTA.

Here we have analyzed two specific regions, namely Region 1 and Region 2, that encompass few of the upstream and intron-2 CpG sites, respectively. While it is worth analyzing methylation at every CpG site in the CSTA locus, our study shows that intron-2 is the site of convergence of estrogen regulation and DNA methylation-dependent silencing. Interestingly, the ERα binding site is located amidst the intron-2 CpGs analyzed in this study. Our global demethylation experiments with MDA-MB-231 and T47D cells revealed the conflict between ERα binding and DNA methylation in region-2. Due to methylation-dependent silencing, MDA-MB-231 cells do not express ERα (Lapidus et al., 1998; Ottaviano et al., 1994) and CSTA (this study). Global demethylation in MDA-MB-231 cells established functional ERα (as revealed by PR induction upon estrogen stimulation), and CSTA mRNA expression. Furthermore, estrogen tends to suppress 5-aza induced CSTA mRNA, resembling estrogen regulation of CSTA in MCF-7 cells. This was possible because demethylation of region 2 CpGs made the intron-2 ERE accessible to ERα. The conflict is also supported by the results from T47D cells. It must be noted that T47D cells show significantly greater level of region 2 methylation than MCF-7 cells, which likely prevents ERα binding to the ERE. This is arguably the reason why despite detectable levels of CSTA and functional ERα in T47D, estrogen does not regulate CSTA. 5-aza not only increased CSTA expression in T47D but also made it amenable to estrogen regulation. Although at this point of time it is not clear why the direction of CSTA regulation in T47D is opposite to that observed in MCF-7, MDA-MB-231 and ZR-75-1 cells, these are enticing evidence that indicate the crucial role of intron-2 in CSTA expression and regulation.

The relationship between DNA methylation and transcription is not a one-way interaction. In a given genomic locus, the transcriptional activity can limit DNA methylation. In Arabidopsis, this is evident from the distribution of methylated and transcriptionally active loci (Zilberman et al., 2007). Pharmacological inhibition of RNA polymerase II induces repressive histone modification, which results in epigenetic silencing. It is possible that repressive histone modification also leads to DNA methylation. Thurman and co-workers studied methylation of transcription factor binding sites and transcription factor abundance in DNAse I hypersensitive sites. They found an inverse correlation between the expression level of a given transcription factor and methylation of the cognate binding site. This suggests a model of “passive DNA methylation” (Thurman et al., 2012). On the other hand, methylation of cytosine residues in CpG dinucleotides prevents binding of transcription factors to their cognate response elements on DNA (Choy et al., 2010; Comb and Goodman, 1990; Miranda and Jones, 2007; Prendergast et al., 1991) thereby interfering with gene expression. An exception to this model is the binding of Sp1 to the methylated cognate site, leading to enhanced gene transcription (Höller et al., 1988). Alternatively, methylated CpG sites attract methyl-cytosine binding proteins (MCBPs) (Hendrich and Bird, 1998). MCBPs in turn recruit histone deacetylases and methylases that cause remodeling and compaction of the local chromatin, and transcriptional shutoff. This is the converse model of methylation-mediated blockade of TF access. In the context of the interaction between transcriptional activity and transcription factor binding, ERα is not an exception. Ung and co-workers have analyzed DNA methylation in relation to ERα expression and binding (Ung et al., 2014). They found an inverse correlation between ERα expression and CpG methylation within ERα binding sites. Methylation of these CpGs was therefore interpreted as being dependent on ERα activity, consistent with the passive model of DNA methylation; more the ERα binding, lesser the methylation. Except for T47D, the cell lines used in this study have patterns of ERα expression and region 2 methylation that is consistent with the passive model. However, that methylation of region 2 in MDA-MB-231 and MDA-MB-453 cells is entirely due to lack of ERα expression, cannot be stated with certainty. Parallelly, results from T47D cells are consistent with the converse model, wherein despite ERα expression and detectable levels of CSTA, estrogen could not significantly modulate CSTA expression. The combined results from MDA-MB-231 and T47D showed that demethylation can restore estrogen regulation at the CSTA locus. We believe that this is most likely due to the restoration of the ERα access to the ERE.

Taken together, the present study shows that CSTA expression in breast cancer cells is inversely related to DNA methylation in the CSTA locus. It explains the loss of CSTA expression in breast tumors. This result may be exploited for predicting CSTA expression or metastatic progression of breast tumors. Furthermore, this study has uncovered an interesting interplay between ERα binding and transcriptional regulation in the CSTA locus. We propose a model in which CSTA expression in breast cancer cells is an integrated result of estrogen regulation and DNA methylation-dependent silencing converging on the intron-2. Our study provides the framework for epigenetic restoration of CSTA expression in breast tumors using pharmacological agents and understanding the role of estrogen in breast tumor invasion and metastasis.

## Acknowledgements

This work was supported by a grant from the Department of Biotechnology (DBT), Govt. of India (Sanction order No. 6242-P90/RGCB/PMD/DBT/ANLM, to A.M.L.). We acknowledge the support from the DBT funded Bioinformatics Facility at IIT Guwahati, and infrastructural support from IIT Guwahati. We thank Mohan C. Manjegowda for his help in ChIP-Seq analysis.

