## Supplementary Table 1 for "Interplay of ERα binding and DNA methylation in the intron-2 determines the expression and estrogen regulation of Cystatin A in breast cancer cells"

**Supplementary table 1:** List of primers.

| Primer name | Sequence | Amplicon length (bp) | Annealing temperature (°C) | Remarks |
|---|---|---|---|---|
| CSTA-F | ATCTGAGGCCAAACCCGCC | 275 | 60 | Used in routine RT-PCR and qRT-PCR |
| CSTA-R | AGCCCGTCAGCTCGTCATC | | | |
| CycA-F | GGGCCGCGTCTCCTTTGAGC | 158 | 60 | Used in routine RT-PCR and qRT-PCR |
| CycA-R | GGCGTGTGAAGTCACCACCC | | | |
| ERα-F | GCCCTACTACCTGGAGAA | 132 | 60 | Used in routine RT-PCR |
| ERα-R | CCCTTGTCATTGGTACTGG | | | |
| CSTA-F | TCCTACTGGATCTCAGCCAC | 369 | 60 | Used in ChIP |
| CSTA-R | GCCCTGTTCTTAGAATAGTGC | | | |
| pS2-F | CATTGCCTCCTCTCTGCTCC | 423 | 60 | Used in ChIP |
| pS2-R | ACTGTTGTCACGGCCAAGCC | | | |
| BST1-F | TAATTTTGATATTATTAGTAAGTTTTGT | 489 | 57 | Used in bisulfite sequencing to amplify Region 2 |
| BST1-R | ATCAACAATCTCCTAAATTTCTAAAA | | | |
| BST2-F | TTTTGTTATTAAATTTTTTTTATTGGATT | 388 | 55 | Used in bisulfite sequencing to amplify Region 1 |
| BST2-R | AAACCCTATTCTTAAAATAATACATAA | | | |

F and R indicate sense and antisense primers respectively.
